# A toolkit for tissue-specific protein degradation in *C. elegans*

**DOI:** 10.1101/104398

**Authors:** Shaohe Wang, Ngang Heok Tang, Pablo Lara-Gonzalez, Bram Prevo, Dhanya K. Cheerambathur, Andrew D. Chisholm, Arshad Desai, Karen Oegema

## Abstract

Proteins essential for embryo production, cell division, and early embryonic events are frequently re-utilized later in embryogenesis, during organismal development, or in the adult. Examining protein function across these different biological contexts requires tissue-specific perturbation. Here, we describe a method that utilizes expression of a fusion between a GFP-targeting nanobody and SOCS-box containing ubiquitin ligase adaptor to target GFP tagged proteins for degradation. When combined with endogenous locus GFP tagging by CRISPR-Cas9 or rescue of a null mutant with a GFP fusion, this approach enables routine and efficient tissue-specific protein ablation. We show that this approach works in multiple tissues—the epidermis, intestine, body wall muscle, sensory neurons, and touch neurons—where it recapitulates expected loss-of-function mutant phenotypes. The transgene toolkit and the strain set described here will complement existing approaches to enable routine analysis of the tissue-specific roles of *C. elegans* proteins.

## INTRODUCTION

Techniques for disrupting protein function in specific tissues or at particular points in development are enabling detailed analysis of developmental mechanisms. To analyze gene function in specific tissues, Cre-LoxP based knockout methods have been established in many model organisms, including C. elegans (Ruijtenberg and Van Den Heuvel, 2015). Tissue-specific CRISPR-Cas9 based gene knockouts and RNAi have also been described in *C. elegans* (Qadota et al., 2007; Shen et al., 2014). However, the utility of DNA/RNA editing approaches can be limited by perdurance of the target protein following excision, which can delay the manifestation of phenotypes.

An alternative approach that circumvents this problem is to directly target proteins for degradation in specific tissues. One method for achieving this is based on transplanting the auxin-induced protein degradation system from plants (Holland et al., 2012; Nishimura et al., 2009). In this system, addition of the small molecule auxin activates a plant-specific F-box protein, TIR1, that serves as a substrate recognition component of an Skp1–Cullin–F-box (SCF) E3 ubiquitin ligase. Active TIR1 targets proteins containing a specific degron sequence. The auxin-inducible degron (AID) system was recently adapted for *C. elegans* (Zhang et al., 2015). However, it would be useful to have a robust genetically-encoded method that does not require a small molecule, as the exposure kinetics and dosage of small molecules in *C. elegans* is limited by barriers such as the cuticle and the eggshell. To this end, a method was developed that takes advantage of an endogenous *C. elegans* protein degradation system. In this approach, target proteins are tagged with a short degron sequence (ZF1), and the SOCS-box adaptor protein ZIF-1, which targets ZF1-containing proteins for proteasomal degradation, is expressed in the target tissue (Armenti et al., 2014). However, a limitation of this system is that ZIF-1 plays an essential role during early embryogenesis (DeRenzo et al., 2003; Reese et al., 2000). Thus, proteins tagged with the ZF1 degron are degraded during embryogenesis as well as in the target tissue, which is problematic for analysis of proteins that function during embryogenesis as well as at later developmental stages.

Here, we develop a new system that combines potent ZIF-1-mediated protein degradation (Armenti et al., 2014) with the previously described deGradFP approach (Caussinus et al., 2012). In deGradFP, a GFP nanobody is fused to a F-box protein to degrade GFP-tagged proteins. Since the originally described F-box adaptor does not work in *C. elegans* (our unpublished observations), we fused the GFP nanobody to ZIF-We show that expression of this fusion enables efficient depletion of GFP-tagged proteins in multiple tissues. In conjunction with GFP-tagging at endogenous loci using CRISPR-Cas9 (Paix et al., 2016) or rescue of null mutants with GFP fusions expressed from transgenes, this approach enables routine protein depletion controlled by the spatial and temporal expression pattern of the promoter driving the GFP-degrading module. We describe a toolkit of transgenes expressing GFP degradation adaptors in different tissues that should facilitate tissue-specific analysis of protein function in *C. elegans*.

## RESULTS

### Epidermal expression of a GFP nanobody::ZIF-1 fusion depletes GFP-tagged proteins that localize to different subcellular locations

To selectively degrade GFP-tagged proteins, we expressed a GFP nanobody::ZIF-1 fusion under tissue-specific promoters (**Fig. 1A**). This fusion protein acts as a GFP-to-ligase adapter that promotes ubiquitination of the GFP-tagged protein by the Cul2 family E3 ligase CUL-2 and subsequent degradation by the proteasome (DeRenzo et al., 2003). We previously showed that epidermis-specific expression of a GFP nanobody::ZIF-1 fusion (*epiDEG;* **Fig. 1A**) led to efficient degradation of an endogenously tagged GFP fusion with the γ-tubulin complex component GIP-2 (Wang et al., 2015). To test whether *epiDEG* can target proteins that localize to different subcellular locations, we crossed the *epiDEG* transgene into strains expressing GFP tagged proteins that localize to: the cytoplasm (transgene encoded GFP::β-tubulin; **Fig. 1B**), apical cell junctions (transgene encoded DLG-1::GFP; **Fig. 1C**), and the nucleus and nuclear envelope (endogenously tagged GFP::MAD-1 (also called MDF-1 in the *C. elegans* literature); **Fig. 1D**). Quantification (**Fig. S1A-C**) revealed a reduction in GFP fluorescence intensity in the larval epidermis for all three markers (DLG-1::GFP, 96%; GFP::MAD-1, 81% GFP::β-tubulin, 58%; **Fig. 1B-D**), whereas signal intensity was unchanged in control tissues (**Fig. 1C-D;** DLG-1::GFP & GFP::MAD-1, GFP::β-tubulin was expressed in the epidermis only). We conclude that the GFP nanobody::ZIF-1 fusion can degrade proteins that localize to different subcellular locations; we note that it remains unclear whether depletion of proteins from the nucleus and cellular junctions occurs by targeting and degradation at these locations or by degrading the protein from the cytoplasm. With the exception of GFP::β-tubulin, which we expect is heavily expressed, reduction by *epiDEG* was consistently greater than 80% (**Fig. 1C-D**; Wang et al., 2015).

**Figure 1.**
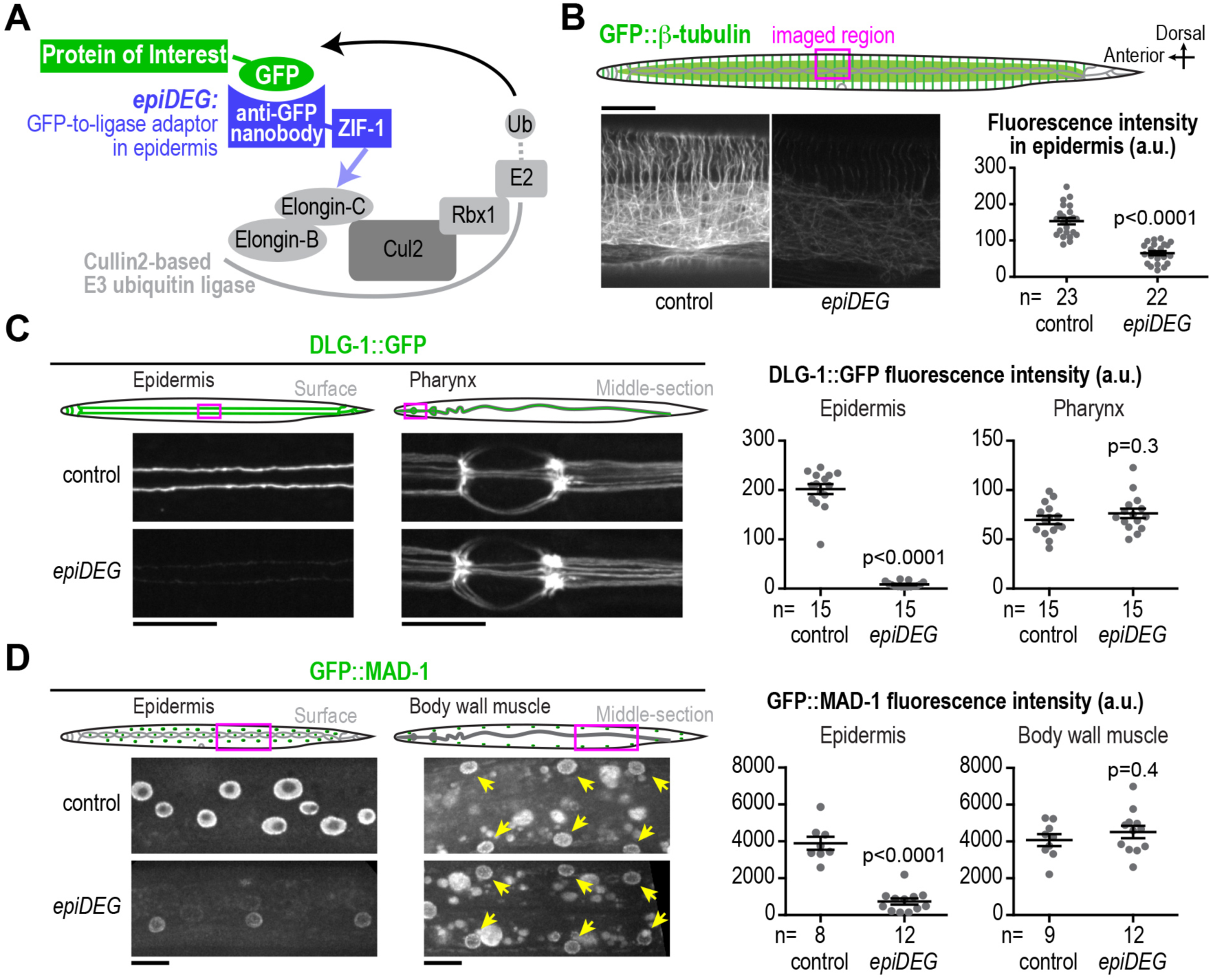
*epiDEG* efficiently degrades GFP-tagged proteins that localize to different subcellular localizations. **(A)** Schematic illustrating the method. (**B**) Top: schematic showing imaged region. Bottom: fluorescence confocal images of L3 stage worms expressing GFP::β-tubulin and plots of GFP::β-tubulin fluorescence intensity. (**C**) Left: schematics and fluorescence confocal images of late L4 stage worms expressing DLG-1::GFP. Right: plots of DLG-1::GFP fluorescence intensity. (**D**) Left: schematics and fluorescence confocal images of L3 stage worms expressing GFP::MAD-1. Right: plots of GFP::MAD -1 fluorescence intensity. n is the number of worms analyzed. Statistics, Student's t-test. p-values are the probability of obtaining the observed results assuming the test group is the same as control. Error bars are SEM. Scale bars, 10 µm.

### GFP-mediated protein degradation is efficient in multiple *C. elegans* tissues

To determine whether the GFP nanobody::ZIF-1 fusion could degrade GFP-tagged proteins in different tissues, we expressed the GFP nanobody::ZIF-1 fusion or ZIF-1 alone (as a control) using promoters that drive expression in the intestine (*intDEG*, P*elt-*2; Fukushige et al., 1998), body wall muscle (*bwmDEG*, P*myo-3*; Fire and Waterston, 1989) and sensory neurons (*senNeuDEG*, P*osm-6*; Collet et al., 1998). The transgenes also included an operon linker (Huang et al., 2001) followed by an mCherry::Histone H2b reporter to allow identification of cells expressing the degradation module (*DEG*) or control transgenes (**Fig. 2A**). To assess the relative function of the degradation module in different tissues, the transgenes were introduced into a background expressing endogenously tagged GFP::MAD-1, which is widely expressed and localizes to nuclei in differentiated tissues throughout development (**Fig. S2**). In all three tested tissues, GFP::MAD-1 signal was eliminated when the GFP nanobody::ZIF-1 fusion, but not ZIF-1 alone, was expressed (**Fig. 2B-D**). Testing *intDEG* with a second endogenously tagged transgene GFP::PP1^GSP-2^ also revealed specific reduction of the intestinal signal to background levels (**Fig. S3**). We conclude that the GFP nanobody::ZIF-1 fusion targets GFP-tagged proteins for degradation in multiple tissues (promoters and targets are summarized in **Table S1**).

**Figure 2.**
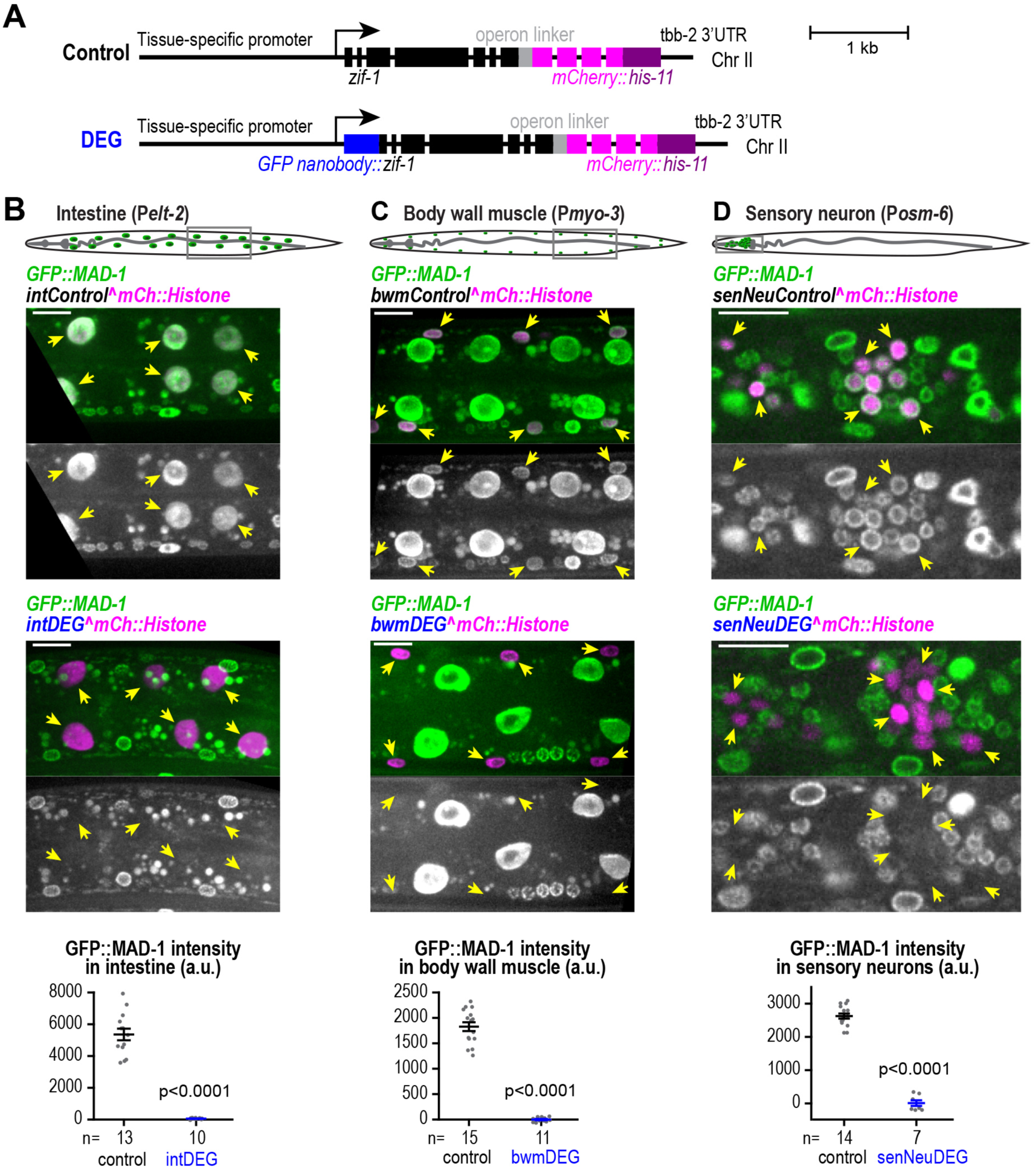
GFP-mediated protein degradation is efficient in multiple *C. elegans* tissues. (**A**) Transgene schematics. (**B-D**) Top: schematics showing imaged region.Middle: fluorescence confocal images (maximum intensity projections in B and C, single z-slice in D) of L3 stage worms expressing GFP::MAD-1. Bottom: plots of GFP: MAD-1 fluorescence intensity. n is the number of worms analyzed. Statistics, Student's t-test. p-values are the probability of obtaining the observed results assuming the test group is the same as control. Error bars are SEM. Scale bars, 10 µm.

### Degradation of GIP-2::GFP in the intestine causes cell division defects and impairs *C. elegans* growth

To determine whether GFP-mediated degradation recapitulated loss-of-function phenotypes, we analyzed embryonic and larval phenotypes in control and *intDEG* worms crossed with endogenously tagged GIP-2::GFP, an essential component of the microtubule-nucleating γ-tubulin complex. During *C. elegans* intestinal differentiation in the embryo, the γ-tubulin complex re-localizes from centrosomes to the apical cell surface (Feldman and Priess, 2012; **Fig. 3A**). Co-expressing *intDEG*, but not the control transgene, eliminated the intestinal GIP-2::GFP signal (**Fig. 3A**).

**Figure 3.**
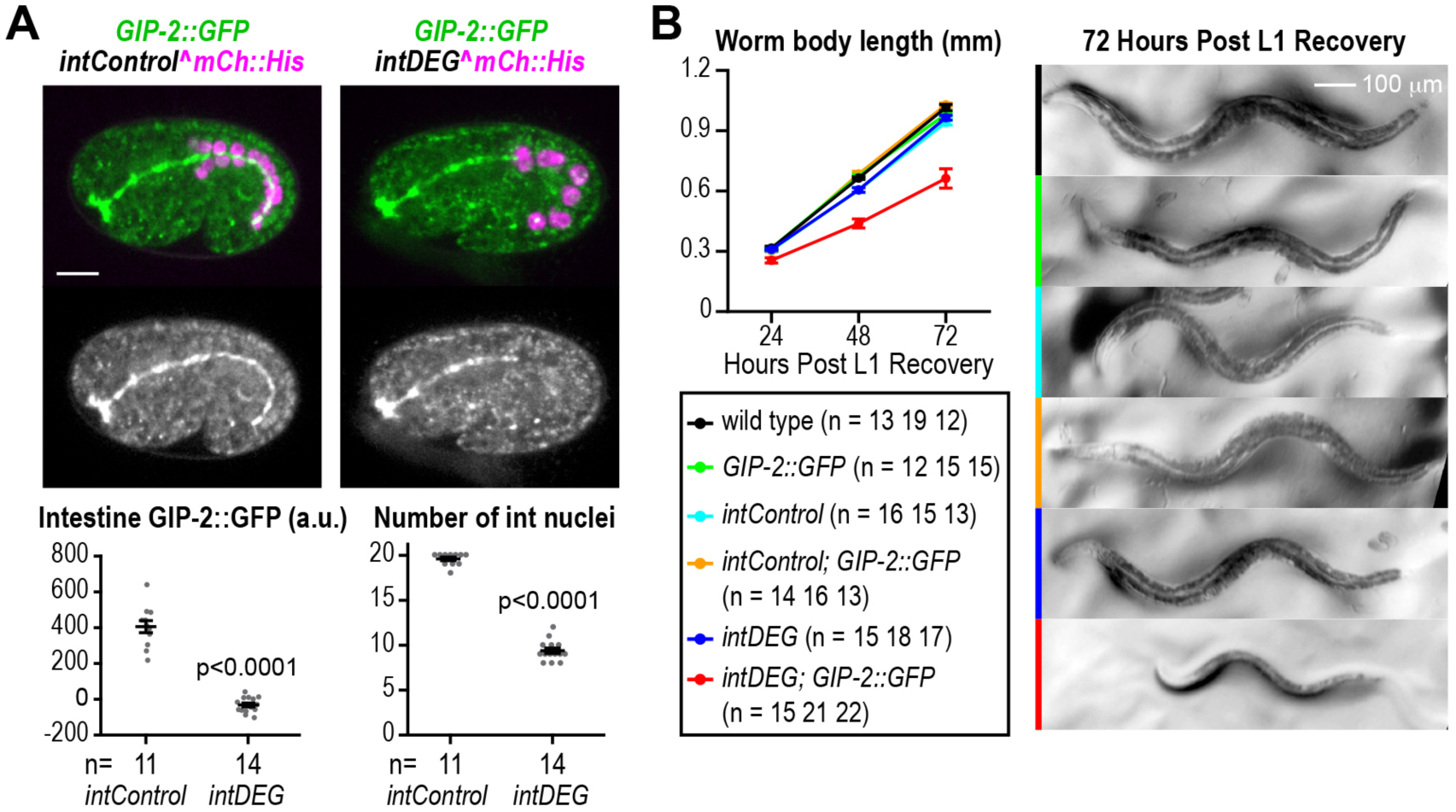
Degradation of GIP-2::GFP in the intestine causes cell division defects and impairs growth. (**A**) Top: maximum intensity projectionsof confocal images of 1.5-fold *C. elegans* embryos.Bottom: plots of GIP-2: GFP fluorescence intensity (*left*) and number of intestinal nuclei (*right*) in 1.5-1.8-fold embryos. n is the number of embryos analyzed. (**B**) Left: plot of body length after recovery from L1 synchronization. Right: representative images of worms 72 hours post recovery. n is the number of worms analyzed at 24, 48 and 72 hours. Statistics, Student's t-test. p-values are the probability of obtaining the observed results assuming the test group is the same as control. Error bars are SEM. Scale bars, 10 µm or as indicated.

The *C. elegans* intestine arises from the E blastomere of the 8-cell embryo (Deppe et al., 1978). The *elt-2* promoter that drives *intDEG* expression turns on at the 2E stage (2-cell intestine) and becomes dominant from the 8E/16E stage (8- to 16-cell intestine) (McGhee et al., 2007). Since the γ-tubulin complex is required for cell division (Hannak et al., 2002; Strome et al., 2001), its inhibition should reduce intestinal cell number. Indeed, in 1.5 to 1.8-fold stage embryos, when control embryos typically have 20 intestinal cells, only 8-10 intestinal nuclei were detected in *intDEG* embryos with endogenously tagged GIP-2::GFP (**Fig. 3A**); larval *intDEG* worms also grew more slowly and reached a smaller adult size than controls (**Fig. 3B**). Thus, intestinal GIP-2::GFP degradation resulted in a tissue-specific cell division defect consistent with loss of the γ-tubulin complex, validating our approach.

### Degradation of GFP::DLK-1 in the touch neurons blocks axon regeneration

To determine whether GFP-mediated protein degradation occurs in the nervous system, we expressed the GFP nanobody::ZIF-1 fusion from a transgene that also included cytoplasmic mKate2 (to visualize axons) using the *mec-18* promoter (*tchNeuDEG*; **Fig. 4A).** To assess efficacy we targeted a GFP fusion with DLK-1 (*D*ual-*L*eucine zipper *K*inase MAPKKK) (K. Noma and Y. Jin, unpublished), which is required to initiate axon regeneration (Hammarlund et al., 2009; Yan et al., 2009). Following laser-induced axotomy in the PLM touch neuron (**Fig. 4B**), GFP::DLK-1 promoted axon regrowth in the presence of endogenous DLK-1, consistent with the known effects of DLK-1 overexpression (Hammarlund et al., 2009; Yan et al., 2009), while a *dlk-1* deletion mutant strongly impaired regrowth (**Fig. 4C, D**). GFP::DLK-1 expression fully rescued the impaired regrowth of the *dlk-1Δ* mutant, and this rescue was abolished by introduction of *tchNeuDEG* (**Fig. 4C-E**). We note that no defects in axon regrowth were detected in *tchNeuDEG* worms with endogenously tagged GIP-2::GFP or DHC-1::GFP (**Fig. S4**), suggesting that GIP-2 and DHC-1 are not required for axon regrowth in PLM touch neurons.

**Figure 4.**
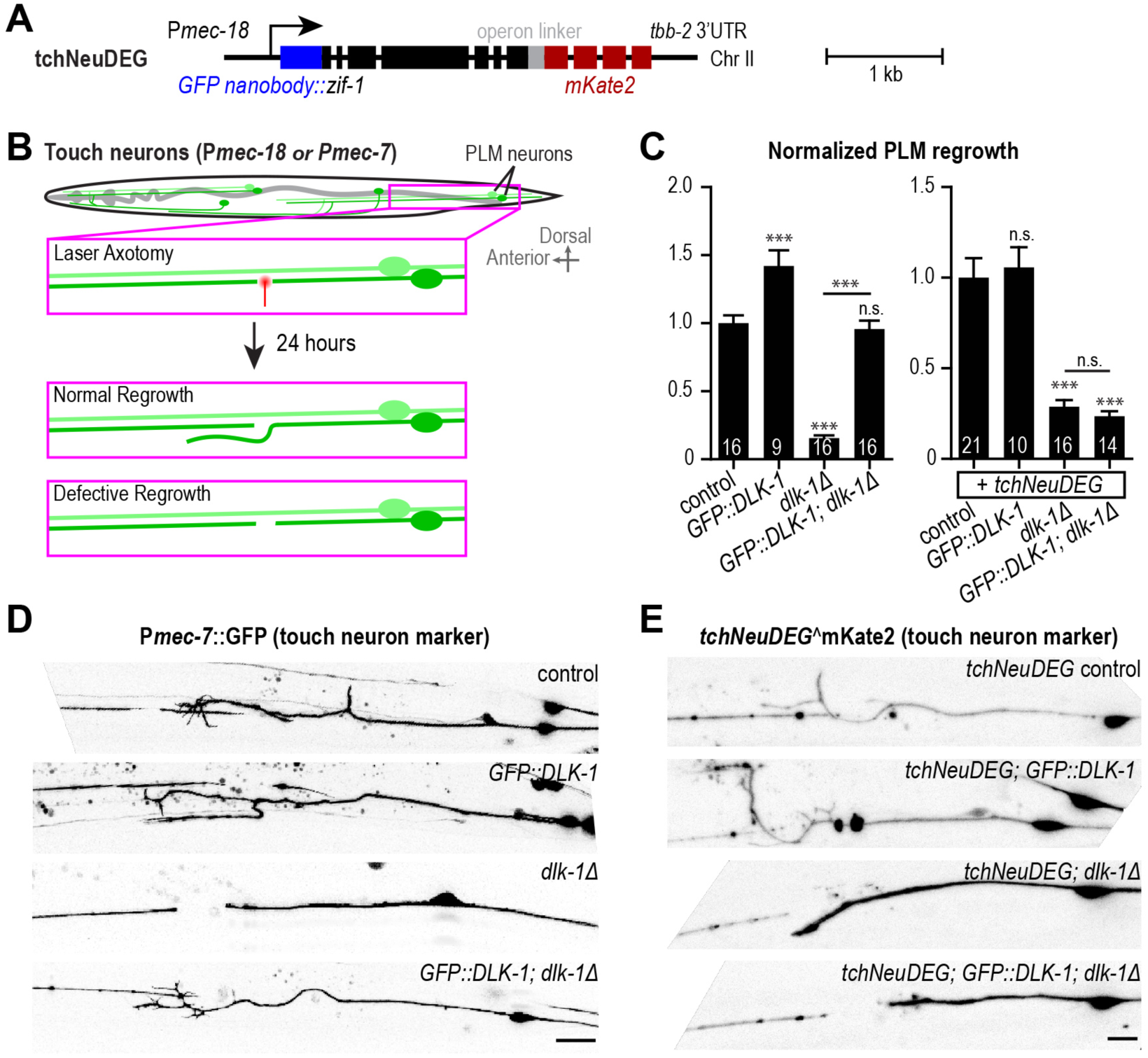
Degradation of GFP::DLK-1 in the touch neurons blocks axon regeneration. (**A**) Transgene schematic. (**B**) Schematic of the axon regeneration assay. (**C**) Plots of normalized touch neuron (PLM) regrowth at 24 hours post laser axotomy. Number in each bar is worms assayed. (**D-E**) Inverted grayscale images of the touch neuron (PLM) axon. Statistics, One-way ANOVA with Bonferroni’s post test. ***: p < 0.001. n.s., not significant. Significance compared to control unless specified by the line. Error bars are SEM. Scale bars, 10 µm.

## DISCUSSION

We describe a robust method for the degradation of GFP fusions that we anticipate will complement existing approaches—genetic locus removal, RNA interference and auxin-mediated degradation—to enable tissue-specific analysis of protein function in *C. elegans*. The utility of this approach is enhanced by the recent development of CRISPR/Cas9-based methods that enable routine GFP tagging of endogenous loci (Dickinson et al., 2013; Paix et al., 2016). We expect that the set of strains we describe here will be a useful resource and will serve as a template for engineering of additional versions that will expand the utility of this strategy.

The strength of the promoter driving the degron cassette, the efficiency of CUL-2-dependent proteasomal degradation in the target tissue, and the expression level of the target will influence the kinetics and penetrance of degradation. At present, we do not know the precise kinetics of in vivo degradation. When the ZF1 degron and ZIF-1 pair was used, target degradation occurred with a half-life of 20-30 minutes (Armenti et al., 2014). We expect the GFP nanobody::ZIF-1 fusion to have comparable performance, but degradation kinetics will need to be measured for each degron/target pair when this information is important for interpreting the phenotype. Whether including sequences targeting the GFP nanobody::ZIF-1 fusion to specific compartments (e.g. the nucleus) would improve degradation efficiency in that compartment also remains to be tested.

The GFP nanobody we use here (vhhGFP4; Rothbauer et al., 2006) recognizes common GFP variants such as EGFP, Venus, YFP, EYFP (Caussinus et al., 2012) and superfolderGFP (S.W., unpublished observation) but does not recognize coral-derived red fluorescent proteins. We have preliminary data that the nanobody also does not recognize mNeonGreen (D.K.C., unpublished observation), a lancelet-derived green fluorescent protein distantly related to Aequorea GFP (Shaner et al., 2013) that exhibits robust fluorescence in *C. elegans* (Dickinson et al., 2015). Thus, red fluorescent proteins are a good choice for marker fusions for phenotypic analysis in the presence of the degron; mNeonGreen could also be used in cases where it is not necessary to use the green channel to monitor degradation of the tagged GFP fusion.

## MATERIALS AND METHODS

### *C. elegans* Strains

*C. elegans* strains are listed in the Supplemental Materials and Methods. All strains were maintained at 20˚C. Transgenic strains were engineered as described (Dickinson et al., 2013; Frøkjær-Jensen et al., 2008). Briefly, GFP was fused to the N-terminus of MAD-1 and PP1^GSP-2^, and the C-terminus of GIP-2 and DHC-1 at their endogenous loci using CRISPR-Cas9 (Dickinson et al., 2013). Transgenes of C-terminally tagged DLG-1::GFP, N-terminally tagged GFP::β-tubulin^TBB-2^ and GFP::DLK-1 were generated using Mos1 transposon mediated single copy insertion (Frøkjær-(Jensen et al., 2008). Constructs and strains will be made available through Addgene and the Caenorhabditis Genetics Center (CGC), respectively.

### Laser Axotomy and Light Microscopy

Laser axotomy was performed as described (Chen et al., 2011). Images in **Figs. 1B** and **S1A** were acquired using an inverted Zeiss Axio Observer Z1 system equipped with AxioVision software, a Yokogawa spinning-disk confocal head (CSU-X1), a 63× 1.40 NA Plan Apochromat lens (Zeiss, Oberkochen, Germany), and a Hamamatsu ORCA-ER camera (Model C4742-95-12ERG, Hamamatsu photonics, Shizuoka, Japan). Images in **Figs. 1C, 1D, 3A** and **S1B** were acquired on the same system using an EMCCD camera (QuantEM:512SC, Photometrics, Tucson, AZ). Images in **Figs. 2**, **S1C-D**, **S2** and **S3** were acquired using a Nikon TE2000-E inverted microscope equipped with Andor iQ2 software, a Yokogawa spinning-disk confocal head (CSU-10), a 60× 1.40 NA Plan Apochromat lens (Nikon, Tokyo, Japan) and an EMCCD camera (iXon DV887ECS-BV, Andor Technology, Belfast, United Kingdom). Images in **Figs. 4D,E and S4** were acquired using Zeiss LSM510 (**Fig. 4D**) and LSM710 (**Figs. 4E** and **S4;** Zeiss Plan Apochromat 63× 1.4 NA oil DIC objective) confocal microscopes controlled by ZEN software (Zeiss). Images in **Fig. 3B** were acquired using the DinoEye eyepiece camera (AM7023B, Dino-Lite, Hsinchu, Taiwan) mounted on a Nikon SMZ800 dissection scope using the DinoXcope software (Dino-Lite).

### Image Analysis

Image analysis was first performed in Fiji (ImageJ) in a semi-automated manner aided by customized macros. Either a box or a line was made inside or across the region of interest to measure raw GFP intensities. Raw measurements were analyzed using customized Python scripts to compute final values. For details, see Supplemental Figures and Materials and Methods.

## ACKNOWLEDGEMENTS

The GFP nanobody vhhGFP4 was cloned from pcDNA3_NSlmb-vhhGFP4, a gift from Markus Affolter (Addgene plasmid # 35579). We thank Kentaro Noma and Yishi Jin for sharing the GFP::DLK-1 transgene prior to publication. We thank the Caenorhabditis Genetics Center for strains and members of the Chisholm lab and the Oegema Desai labs for helpful discussions.

## COMPETING INTERESTS

The authors declare no competing interests.

## AUTHOR CONTRIBUTIONS

S.W., N.H.T, A.D.C., A.D. and K.O. designed the experiments. S.W., N.H.T., P. L-G., B.P. and D.K.C performed the experiments. S.W. and N.H.T. performed data analysis. S.W., A.D. and K.O. wrote the paper with input from all authors.

## FUNDING

This work was supported by the National Institutes of Health [GM074207 to K.O., NS093588 to A.D.C.]. A.D. and K.O. receive salary and other support from the Ludwig Institute for Cancer Research.

